# DeepTracer ID: De Novo Protein Identification from Cryo-EM Maps

**DOI:** 10.1101/2022.06.03.494766

**Authors:** Luca Chang, Fengbin Wang, Kiernan Connolly, Hanze Meng, Zhangli Su, Virginija Cvirkaite-Krupovic, Mart Krupovic, Edward H. Egelman, Dong Si

**Affiliations:** Division of Computing and Software Systems, University of Washington Bothell, Bothell, WA 98011; Department of Biochemistry and Molecular Genetics, University of Virginia School of Medicine, Charlottesville, VA 22903; Department of Genetics, University of Alabama at Birmingham, Birmingham, AL 35233; Institut Pasteur, Université de Paris, CNRS UMR6047, Archaeal Virology Unit, 75015 Paris, France

## Abstract

The recent revolution in cryo-electron microscopy has made it possible to determine macromolecular structures directly from cell extracts. However, identifying the correct protein from the cryo-EM map is still challenging and often needs additional sequence information from other techniques, such as tandem mass spectrometry and/or bioinformatics. Here, we present DeepTracer-ID, a server-based approach to identify the candidate protein in a user-provided organism *de novo* from a cryo-EM map, without the need for additional information. Our method first uses DeepTracer to generate a protein backbone model that best represents the cryo-EM map, and this model is then searched against the library of AlphaFold2 predictions for all proteins in the given organism. This method is highly accurate and robust: in all 13 experimental maps tested blindly, DeepTracer-ID identified the correct proteins as the top candidates. Eight of the maps were of known structures, while the other five unpublished maps were validated by prior protein annotation and careful inspection of the model refined into the map. The program also showed promising results for both homomeric and heteromeric protein complexes. This platform is possible because of the recent breakthroughs in large-scale protein 3D structure prediction.

**Statement of Significance:** While it has now become routine for cryo-EM maps of proteins to reach a near-atomic resolution, potentially allowing for reliable atomic models to be built, there are a growing number of instances where the protein identity may not be known. Without knowing the protein sequence, it is impossible to build an atomic model. DeepTracer-ID is a server-based approach to surmount this problem by identifying the proteins in a given organism that are found in the cryo-EM map. A free web service for global academic access is provided.

## Introduction

Over the past nine years, there has been a resolution revolution in cryo-electron microscopy (cryo-EM), with exponential growth in the number of determined atomic structures per year^1-3^. Unlike X-ray crystallography or NMR which often require a large amount of sample, a high concentration and a high degree of sample purity, using cryo-EM it is possible to capture structures of macromolecular complexes under conditions closer to the *in situ* environments with fewer restraints on volume, sample purity and concentration^4-6^. With recent software developments^7,8^, cryo-EM can routinely sort out different conformational states of single macromolecules^9^, and even unrelated complexes within the same dataset^10,11^. When a cryo-EM map of unknown components reaches ∼4.2 Å resolution or better where β-sheets are well resolved, one can typically trace the Cα backbone directly from the map. The remaining challenge is identifying the correct protein sequence from the organism’s genome.

A typical workflow that has been used to address this problem is first to detect what proteins are present in the sample by tandem mass spectrometry, then threading the sequences of detected proteins into the cryo-EM map to identify the best match^12-14^. However, this approach relies on the target protein being detected by MS/MS, which is often not the case due to the lack of digestion sites^15^, a high degree of post-translational modifications^16^, etc. Or on the contrast, thousands of proteins can be detected if the target of interest was purified directly from cell extracts. Under such circumstances, possible sequence targets have to be selected based on the structural features of the map. Then each sequence must be built into the map to identify the best match by trial and error.

A second approach is using model building tools, such as DeepTracer^17^, to perform protein sequence prediction based on an input cryo-EM map. Then, the predicted sequence is used for BLAST analysis to identify potential hits in the selected organism^18^. This approach is greatly limited by the accuracy of the backbone trace and the quality of side chain densities of the input cryo-EM map, and the results may not be straightforward to interpret.

Another approach to identifying unknown proteins is to trace the backbone from the cryo-EM map, and then find the best matches against the experimentally determined structures in the Protein Data Bank. One example of this approach is *FindMySequence*^19^. However, this method is likely to be limited to detecting proteins with known structure or those closely similar to structurally characterized proteins. Fortunately, with recent breakthroughs in protein 3D structure prediction by AlphaFold2, it is possible to obtain accurate protein structure predictions even when no homologous structure is available^20^.

Here, we describe a *de novo* protein identification approach, DeepTracer-ID, for cryo-EM maps with better than 4.2 Å resolution. Our program first automatically generates a backbone tracing of the cryo-EM map using DeepTracer. Then, the AlphaFold2 predicted models are aligned to the DeepTracer backbone using three different algorithms, PyMOL-align^21^, PyMOL-cealign^22^ and FATCAT^23^. On a benchmark set of 11 experimental segmented cryo-EM maps, including three previously unsolved structures, we show that our program, DeepTracer-ID, can identify the correct protein and their closely related homologs in a pool of AlphaFold2 predicted models. Furthermore, we show that DeepTracer-ID can detect the correct protein from unsegmented maps made of homopolymers, and also identify large components from maps of massive complexes.

## Methods

### Pipeline inputs of DeepTracer-ID

Two inputs are needed to use the pipeline: (1) a cryo-EM map, segmented to correspond to a single protein subunit is preferred but not necessary; and (2) A pre-calculated AlphaFold2 protein library to search against (Fig. 1). The input cryo-EM map is used to generate a 3D model trace by DeepTracer^17^. In a rare case when the cryo-EM map does not contain a single α-helix, there is an ambiguity in the absolute hand of the cryo-EM map^24^. In this case, the users are encouraged to mirror the input map and run DeepTracer-ID again for the best results. The proteomes from a number of model organisms and pathogens important to global health, such as *H. sapiens, E. coli, A. thaliana, C. elegans, C. jejuni, S. pneumoniae*, etc., have been already pre-calculated^25^, and for those organisms an AlphaFold2 protein library is not required from the user side. For the other organisms, the user can either generate their AlphaFold2 library locally or use an online service such as ColabFold^26^.

**Fig. 1.**
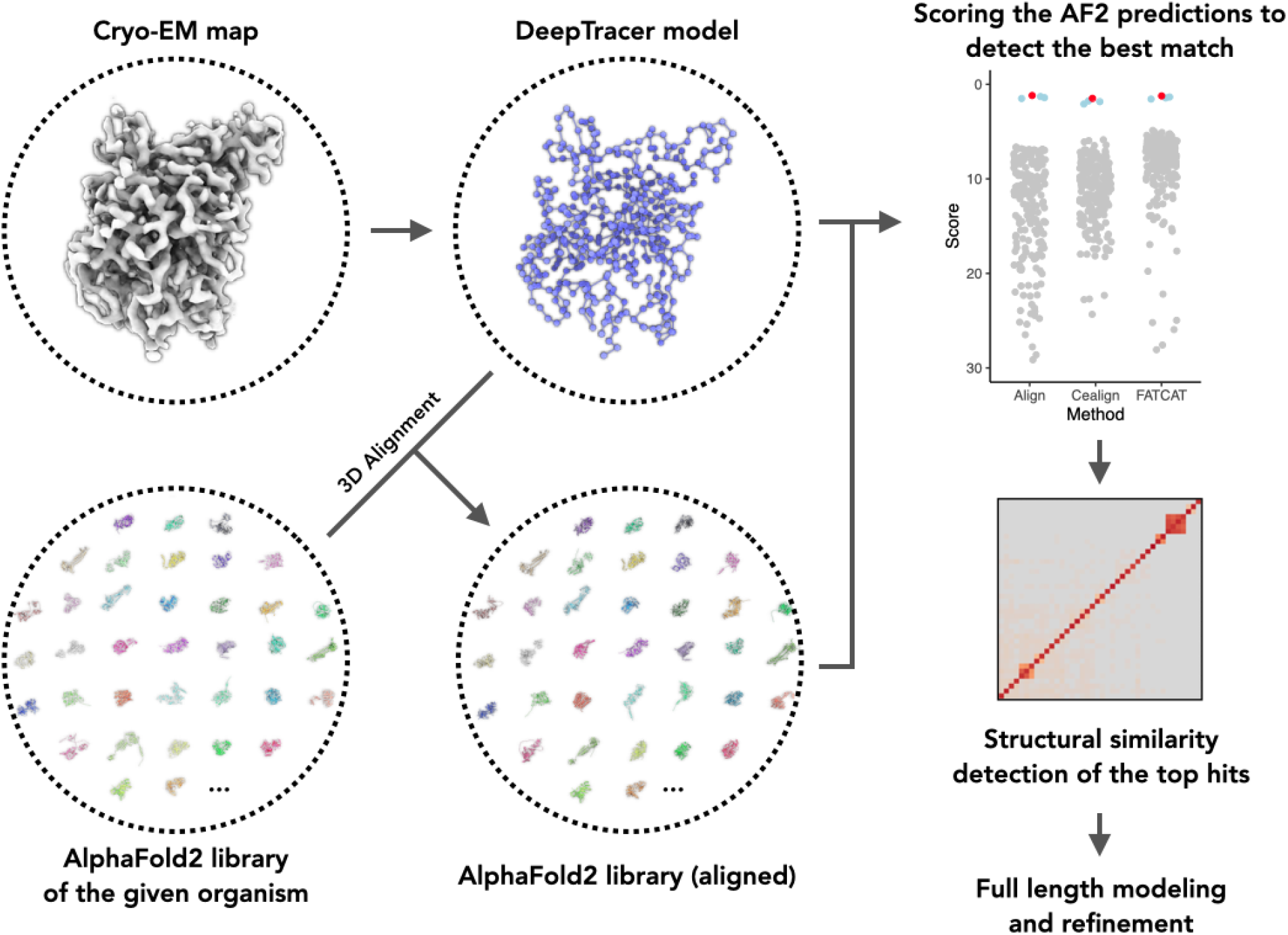
DeepTracer-ID: *de novo* protein identification from cryo-EM density maps. Starting with a cryo-EM map and a given organism, DeepTracer-ID identifies the protein in three steps: (1) Initial model is generated from the cryo-EM map using deep learning based DeepTracer; (2) Align the AlphaFold2 prediction library (user provided or server pre-calculated) to the initial model; (3) Score each aligned AlphaFold2 prediction and subsequent analysis of the top hits.

### 3D alignment of AlphaFold2 predictions to map-traced model

We use three different approaches, PyMOL-align^21^, PyMOL-cealign^22^ and FATCAT^23^, to align the AlphaFold2 predicted structures to the backbone model traced directly from the cryo-EM map (Fig. 1). The aligned AlphaFold2 predictions are saved separately for subsequent RMSD analysis. Three different alignment algorithms are used, as we expect they have different advantages and disadvantages depending on the input cryo-EM map. PyMOL-align considers both similarities in sequence and structure, and this is the default alignment option due to its overall most robust performance (see below). PyMOL-cealign mostly weights on structural similarity, so it may be more useful when the protein side chain densities are not well resolved in the cryo-EM map. FATCAT has a flexible alignment feature so it may be able to minimize some of the errors introduced from the AlphaFold2 prediction, and it may be more useful for smaller proteins, or proteins that rely on the local environment to form an ordered 3D structure.

### Protein identification scoring matrix

Each AlphaFold2 prediction is scored with two factors: (1) the RMSD to the DeepTracer model and (2) the percentage of residues in the DeepTracer model that can be aligned to the corresponding AlphaFold2 prediction (Eq. 1). The RMSD calculation between the AlphaFold2 prediction and the DeepTracer model is done with the methods previously implemented in DeepTracer^17^. The length of the DeepTracer model is *l*_*DT*_and the length of matching residues in the superposed AlphaFold2 prediction is *l*_*aligned*_. The AlphaFold2 predictions are then listed from lowest to highest based on this score, and the correct protein is expected to be detected within the AlphaFold2 predictions with lowest score:

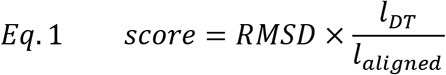

Using this approach, we expect to not only detect the correct protein, but also other isoforms or homologs that are structurally similar to the correct protein. Therefore, after scoring all AlphaFold2 predictions, we use the DaliLite v5^27^ to perform an all-to-all analysis on the top hits.

## Results

### Protein identification from segmented cryo-EM maps

We tested our method on a benchmark set of eight segmented cryo-EM maps from proteins that were known. Those proteins range in length from 224 to 838 residues and the maps have a resolution ranging from 2.6 to 3.9 Å. Initially, five cryo-EM maps were selected as test sets to optimize our approach. The selection criteria considered the most likely pitfalls for this approach, and we tried to include cases where: (1) the map lacks secondary structure and has multiple ligands bound; (2) only a portion of the map reaches high resolution; (3) the corresponding protein has multiple domains, and their relative orientation is unlikely to be accurately predicted by AlphaFold2. In all five cases, despite these existing possible pitfalls, DeepTracer-ID was able to identify the correct protein as the top hit (Fig. 2). We then moved on to test three more EM maps from eukaryotes with larger genomes, two from *H. sapiens a*nd one from *A. thaliana*. DeepTracer-ID could also identify the correct protein as the top hit, except for one case. In that case, the correct protein was ranked 3rd, while the top 32 hits are all structurally related homologs (Figs. 2&3).

**Fig. 2.**
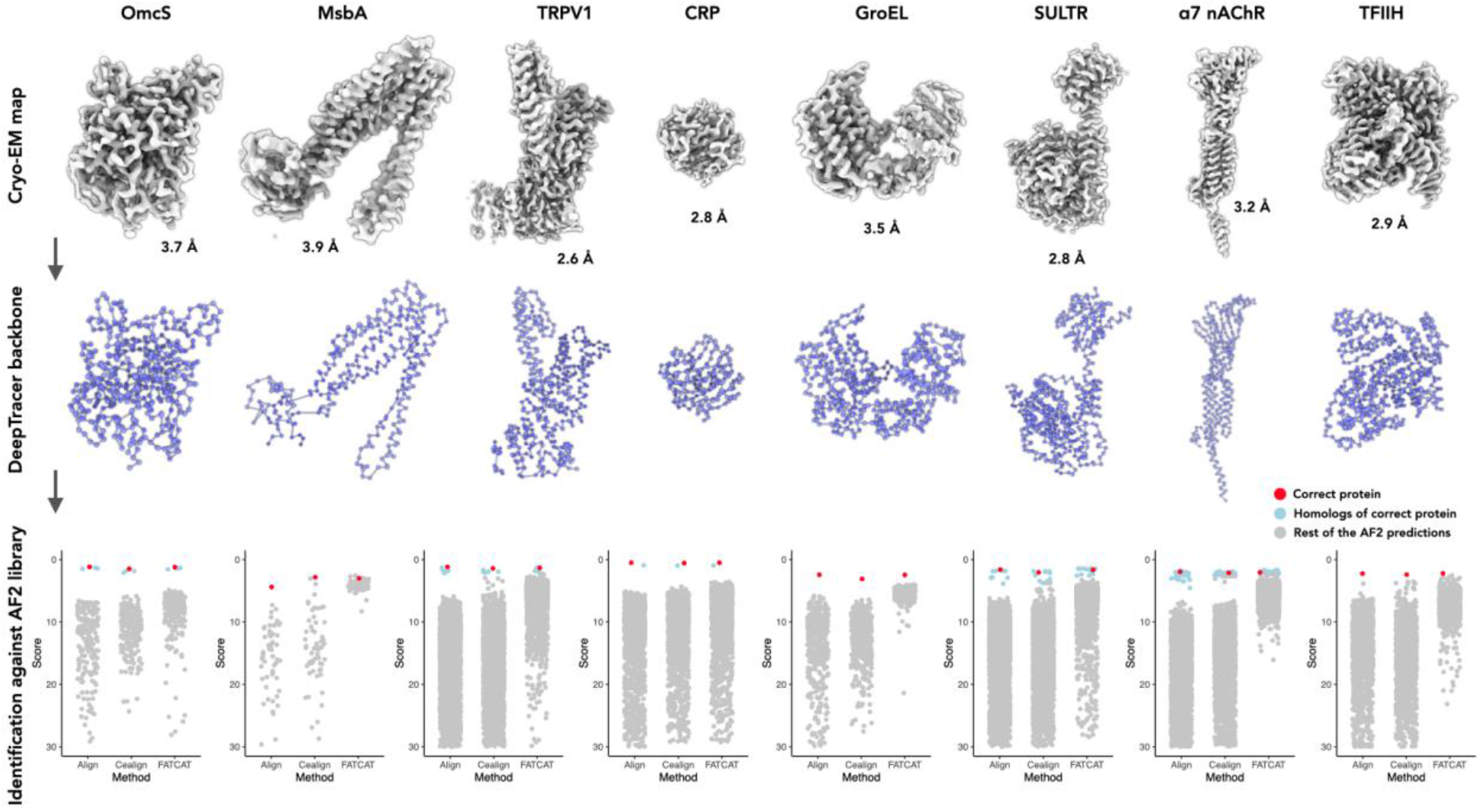
Protein identification from eight benchmark cryo-EM maps. Top, cryo-EM maps used for *de novo* protein identification. Their reported resolution is labeled. Middle, the Cα backbone of the model generated by DeepTracer, according to the maps on top. Bottom, the scores of corresponding AF2 predictions are displayed in scatter plots. The results from all three different alignment algorithms are shown. The correct protein is shown in red dot, the proteins with significant structural similarity to the correct protein are labeled in blue dot, and the rest of AF2 predictions are shown in grey dots. The size of AF2 library and the corresponding organism for the eight benchmark datasets are: OmcS (*G. sulfurreducens PCA*, N=226), MsbA (*E. coli BL21-DE3*, N=65), TRPV1 (*R. norvegicus*, N=3679), CRP (*H. sapiens*, N=1419), GroEL (*E. coli K12*, N=413), SULTR (*A. thaliana*, N=3850), α7 nAChR (*H. sapiens*, N=2847), and TFIIH (*H. sapiens*, N=1347).

**Fig. 3.**
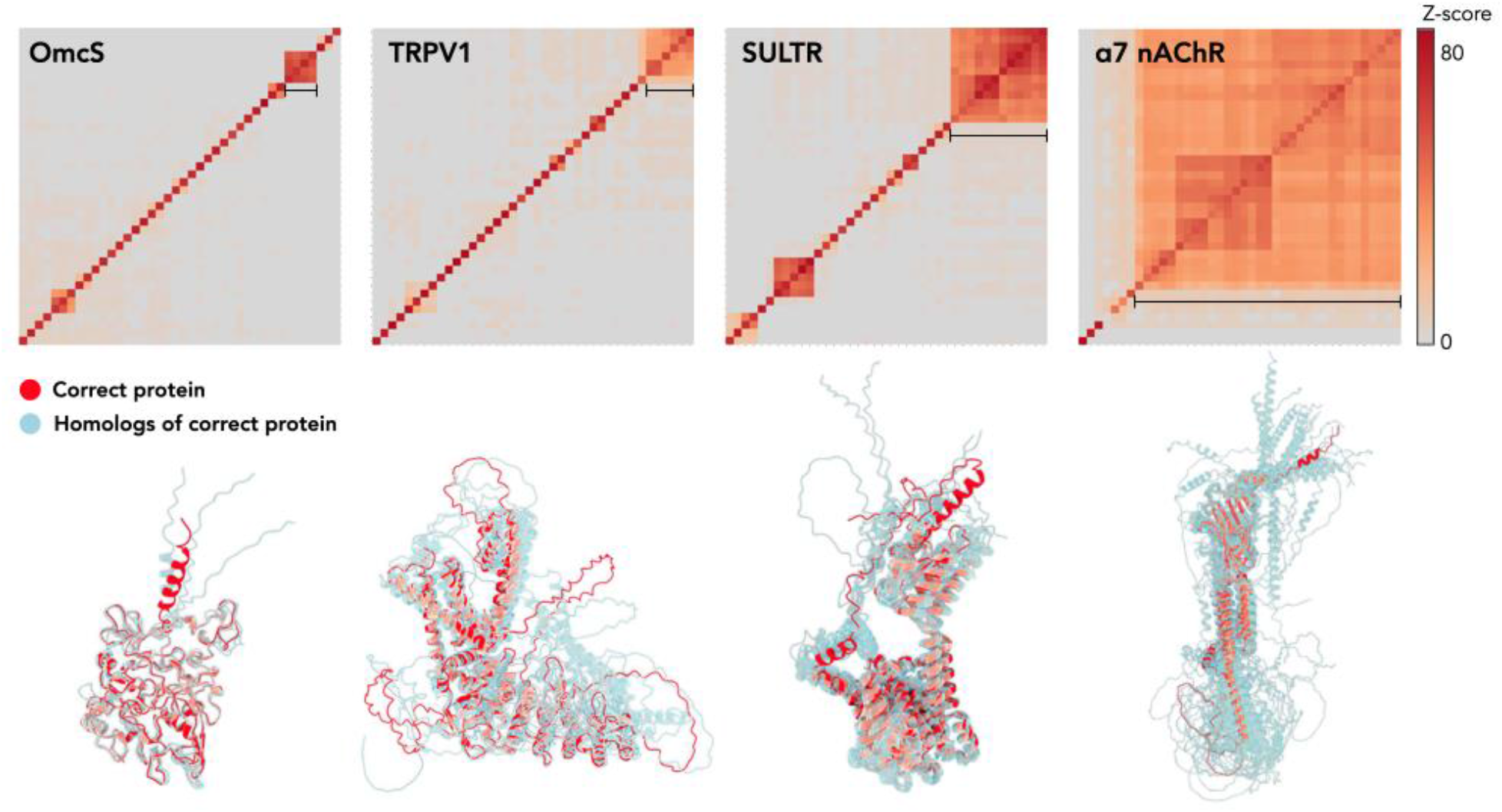
DALI all-to-all analysis of top 40 scoring AF2 predictions. Top, DALI all-to-all analysis of four representative benchmark datasets shown in Fig. 2. The matrix is based on the pairwise Z-score comparisons calculated using the DALI server. The color scale on the right indicates the corresponding Z-scores. Bottom, the structures clustered (indicated by black line) in the top matrix are shown. The correct protein is colored in red. The remaining proteins are colored in transparent cyan and aligned to the correct protein.

Among the proteins in the benchmark set, OmcS^14^ has six hemes as ligands, and TFIIH^28^ has a bound dsDNA. Interestingly, our method detected the correct protein using cryo-EM maps in the presence or absence of ligand/DNA densities (supp Fig. 1), suggesting that a small portion of non-protein density does not diminish the accuracy of our method. Importantly, although OmcS lacks pronounced secondary structure elements, DeepTracer correctly placed Cα atoms into the map, when the resolution was better than 4.2 Å. Therefore, a successful identification primarily relies on the accuracy of AlphaFold2 predictions.

### Closely related structural homologs of the correct protein

If the correct protein has structural homologs/isoforms, we expect both the correct protein and its homologs/isoforms to be detected using our method. This is inevitable because the correct protein and its homologs/isoforms are structurally similar. In addition, model errors should be expected in both Cα tracing in DeepTracer and AlphaFold2 prediction. As a result, the correct protein and its homologs cannot be distinguished at this step. Therefore, a subsequent full-length model building for all top hits is required to determine which protein best fits into the cryo-EM map. This is now a routine protocol for *de novo* model building of an unknown protein^14,15,29^. To visualize possible structural homologs of the top hits, we implemented DALI all-to-all analysis^27^ into this pipeline. This additional DALI analysis will not only detect potential sequence ambiguity, but also reveal possible biologically interesting similarities that were not noticed. As expected, in several cases, our method detected a number of structural homologs of the correct protein (Fig. 3). These can be confidently reconciled by subsequent full-length modeling, without the need for additional data.

### De novo protein identification of unknown cryo-EM maps

As a proof of principle, we then applied our method to three segmented cryo-EM maps with moderate resolutions from 3.4-3.9 Å of unknown proteins. The first protein is an archaeal flagellin from *Aeropyrum pernix*, with a long N-terminal helix and a C-terminal globular domain^30^. Without additional information on how this flagellin packs into a filament, it is challenging for AlphaFold2 to accurately predict how the two domains are oriented with respect to each other. The other two proteins are the two major capsid proteins (MCP) of Acidianus filamentous virus 6 (AFV6), a virus infecting a hyperthermophilic and acidophilic archaeon^31^. Similar to SIFV^32^, the two capsid proteins wrap around A-form DNA. Without the information about both the helical packing and protein-DNA interactions, AlphaFold2 will not be able to predict the entire structure accurately. Also, archaeal and archaeal virus genomes are generally much more sparsely sampled than those from bacteria and eukaryotes and the viruses that infect them. Therefore, fewer homologous sequences are available for AlphaFold2 to use in the prediction.

With all those potential pitfalls, our method still successfully determined the correct sequences (Fig. 4). In all three cases, only 45%-71% of the AlphaFold2 predicted models could be satisfactorily aligned to the backbone traced from the cryo-EM maps. And yet, such coverage is sufficient for our method to detect the correct protein. The full-length models were subsequently built by DeepTracer and real-space refined by PHENIX^33^. The statistics of the deposited models are listed in Supp. Table 1.

**Fig. 4.**
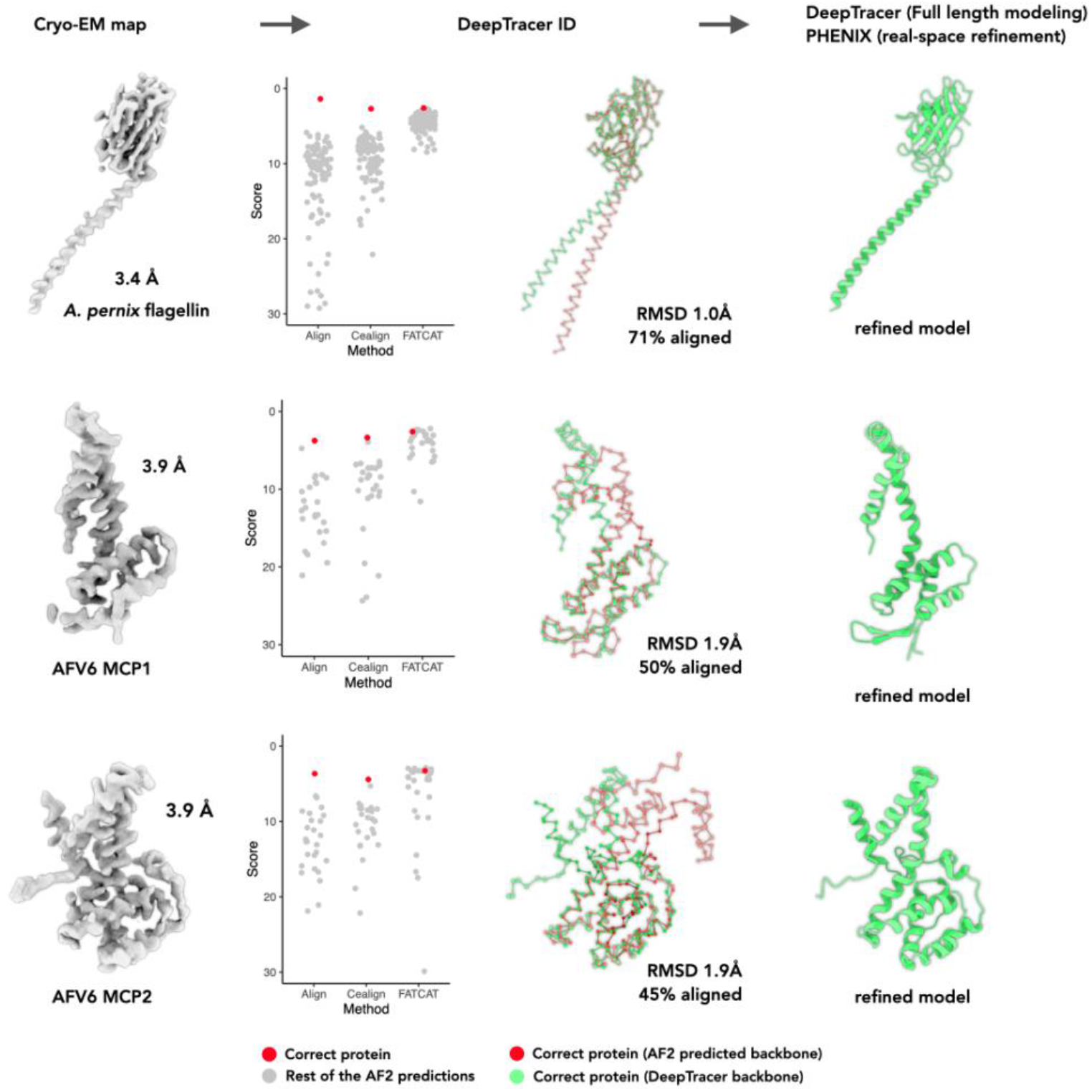
Protein identification from *A. pernix* and archaeal virus AFV6. Far left, cryo-EM maps used for *de novo* protein identification. Left, the scores of corresponding AF2 predictions are displayed in scatter plots. The correct protein is shown in red dot and the remaining AF2 predictions are shown in grey dots. The size of AF2 library and the corresponding organism for the three unpublished datasets are: *A. pernix* flagellin (N=104), AFV6 MCP1 (N=28) and AFV6 MCP2 (N=28). Right, the alignment between DeepTracer model (green) and AF2 prediction of the correct protein (red). The alignment RMSD and percentage of aligned area are also labeled. Far right, the final model after DeepTracer full-length modeling and PHENIX real-space refinement.

### Structural knowledge mined from large and complicated maps

The next question was whether the approach could extract useful information from a more complicated map. We arbitrarily created two tiers for the maps based on their complexity: the ‘easy’ tier maps contain multiple copies of the same protein subunit; the ‘difficult’ tier contains complex maps that are built from multiple components. We tested our method with two very different maps in the easy tier: the dimeric map of human thyroglobulin with two protein copies and the filament map of an *A. pernix* flagellum with ∼50 flagellin copies. Our method successfully determined the correct protein in both cases (Fig. 5). Interestingly, the gap between the correct protein and the protein with the 2nd best score is comparable to the results from segmented maps. This is presumably because the alignment between AlphaFold2 predictions and DeepTracer models is insensitive to the number of protein copies present in the cryo-EM map. In addition, DeepTracer assigns residues to every corner of the map, thus the subsequent alignment is not affected much by whether the backbone corresponding to a single subunit is successfully traced or not.

**Fig. 5.**
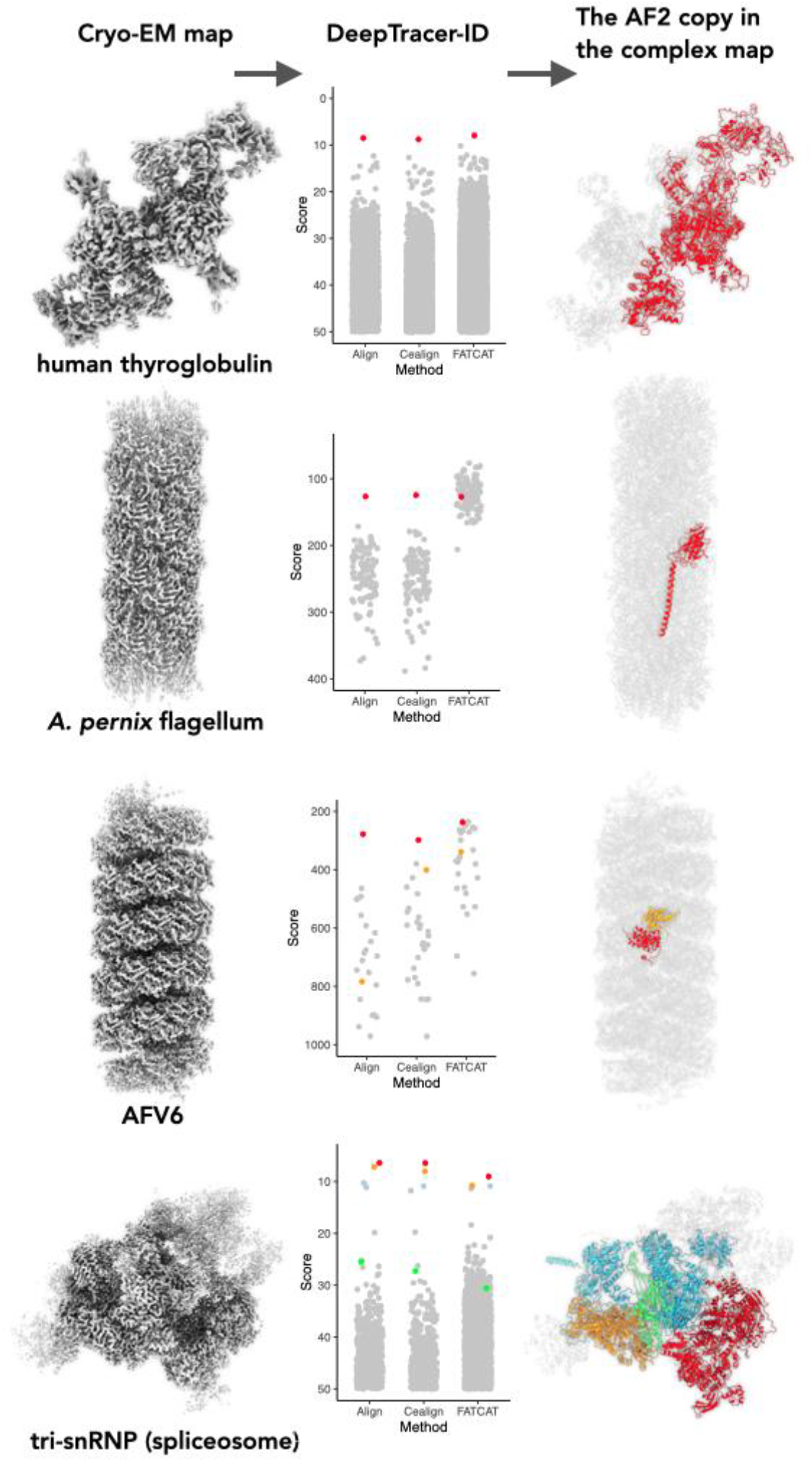
Protein identification from protein complex maps. Left, cryo-EM maps used for *de novo* protein identification. Middle, the scores of corresponding AF2 predictions are displayed in scatter plots. The correct protein is shown in colored dots and the remaining AF2 predictions are shown in grey dots. The size of AF2 library and the corresponding organism for the four datasets are: *H. sapiens* thyroglobulin (N=22742), *A. pernix* flagellin (N=104), and *H. sapiens* tri-snRNP complex (N=23391). Right, the identified protein(s) are colored in the same theme as shown in the middle, the rest of the complex is colored in grey.

Next, we were curious whether DeepTracer-ID could extract any useful information from complicated cryo-EM maps containing multiple components. To this end, we tested two maps (Fig. 5). The first map is that of the filamentous virus AFV6, with ∼50 copies of MCP1, ∼50 copies of MCP2, and ∼600 bp of double-stranded A-form DNA. Interestingly, our method could still detect one of the capsid proteins, MCP2. However, the other capsid protein did not rank high using the default PyMOL-align algorithm. It ranked 3rd using PyMOL-cealign, suggesting that sequence information may not be helpful in a highly complicated map. The second map tested is the tri-snRNP part of the spliceosome complex composed of ∼30 different components. Not surprisingly, our method detected the largest four proteins in the complex, as they are expected to have a higher *l*_*aligned*_ in Eq. 1, leading to a better score. This suggests that DeepTracer-ID can identify the major components from large complex maps, thereby rendering the subsequent map segmentation and smaller protein identification much easier.

## Discussion

We present here a Cryo-EM pipeline, DeepTracer-ID, that can robustly identify the protein components directly from cryo-EM maps of a given organism, without the need for other experimental data. Furthermore, this approach does not require very high accuracy from AlphaFold2 predicted libraries. As long as ∼45% of the AlphaFold2 prediction reasonably matches the cryo-EM structure, this pipeline can identify the correct protein. As a proof of principle, we successfully identified three proteins within large filamentous complexes directly isolated from natural sources using this pipeline.

The remaining challenge for this pipeline is working with a map of tiny protein. Due to the extremely high signal-to-noise level in cryo-EM, it is currently almost impossible to reach a near-atomic resolution for a small monomeric protein (<100 amino acids). However, such small proteins frequently exist in larger complexes, such as conjugation pili^34,35^ and ciliary doublets^29^. In such cases, the DeepTracer model can probably be structurally aligned to many different AlphaFold2 predictions reasonably well. At the same time, sequence information may no longer be beneficial due to the limited length. For example, we tested a virus capsid protein with only ∼90 amino acids in a segmented cryo-EM map (*Wang et al. Cell 2022 in press*, supp Fig. 2). The protein has the N-terminal ∼60 residues cleaved, a helix-turn-helix structure, glycosylation in the middle, and no cleavage site for trypsin. As a result, we found that this set of features rendered the initial sequence-based alignment method unreliable, while the structure-based PyMOL-cealign remained valid. FATCAT can increase the score of the correct protein by enabling flexible fitting, but it improves the scores for all other proteins at the same time. We will consider introducing other initial alignment methods in the future, such as Chimera matchmaker^36^ or different flexible fitting algorithms.

This approach could be broadly applicable to many scenarios other than identifying an unknown protein in a cryo-EM map resulting from contaminants. Using this method, it is possible to investigate large complexes enriched in a cell extract. The protein of interest can be directly identified when the resolution is better than 4.2 Å, regardless of post-translational modifications and the lack of proteomics data. It may also be utilized to do a cryo-EM ‘pull-down’ assay, where instead of a long list of candidates from mass spectrometry, the outputs now are protein 3D structures fitted to the maps.

## Data availability

The DeepTracer-ID described here is free for academic use, available at https://deeptracer.uw.edu/home. The atomic models and cryo-EM volumes have been deposited in the Protein Data Bank and the Electron Microscopy Data Bank (*A. pernix* flagellum, 7TXI and EMD-26158; AFV6, 7TXJ and EMD-26159). Other data are available from the corresponding authors upon reasonable request.

## Author Contributions

F.W., E.H.E., and D.S. initiated and developed the project. F.W. performed microscopy and image analysis. V.C-K. and M.K. cultured the viruses and archaeal cells and performed sample preparation. L.C. and K.C. wrote the code and built the web service of DeepTracer-ID. F.W. prepared the benchmark datasets for algorism testing. F.W. and Z.S. analyzed data and developed graphical representations. F.W., E.H.E., and D.S. wrote the manuscript with input from all authors.

## Acknowledgments

The Cryo-EM imaging was done at the Molecular Electron Microscopy Core Facility at the University of Virginia, which is supported by the School of Medicine and built with NIH grant G20-RR31199. This work was supported by NIH Grant GM122510 (E.H.E.), K99GM138756 (F.W.), NSF grant 2030381 (D.S.), the SRCP Seed Grant at the University of Washington Bothell (D.S) and l’Agence Nationale de la Recherche grant ANR-21-CE11-0001-01 (M.K.).

## Materials and Methods

**Table S1.** Cryo-EM and Refinement Statistics of *A. pernix* flagellum and AFV6 filaments.

| Parameter | *A. pernix* flagellum | AFV6 |
| --- | --- | --- |
| **Data collection and processing** |  |  |
| Voltage (kV) | 300 | 300 |
| Electron exposure (e<sup>-</sup> Å<sup>-2</sup>) | 50 | 50 |
| Pixel size (Å) | 1.08 | 1.4 |
| Particle images (n) | 59,338 | 78,141 |
| Shift (pixel) | 8 | 10 |
| **Helical symmetry** |  |  |
| Point group | C1 | C1 |
| Helical rise (Å) | 5.52 | 5.75 |
| Helical twist (°) | 108.0 | 38.46 |
| **Map resolution (Å)** |  |  |
| Map:map FSC (0.143) | 3.5 | 3.9 |
| Model:map FSC (0.38) | 3.7 | 4.2 |
| d<sub>99</sub> | 3.9 | 4.1 |
| **Refinement and Model validation** |  |  |
| Ramachandran Favored (%) | 93.2 | 93.1 |
| Ramachandran Outliers (%) | 0.5 | 0.6 |
| RSCC | 0.82 | 0.85 |
| Clashscore | 9.6 | 12.6 |
| Bonds RMSD, length (Å) | 0.004 | 0.006 |
| Bonds RMSD, angles (°) | 0.732 | 0.781 |
| **Deposition ID** |  |  |
| PDB (model) | 7TXI | 7TXJ |
| EMDB (map) | EMD-26158 | EMD-26159 |

**Fig. S1.**
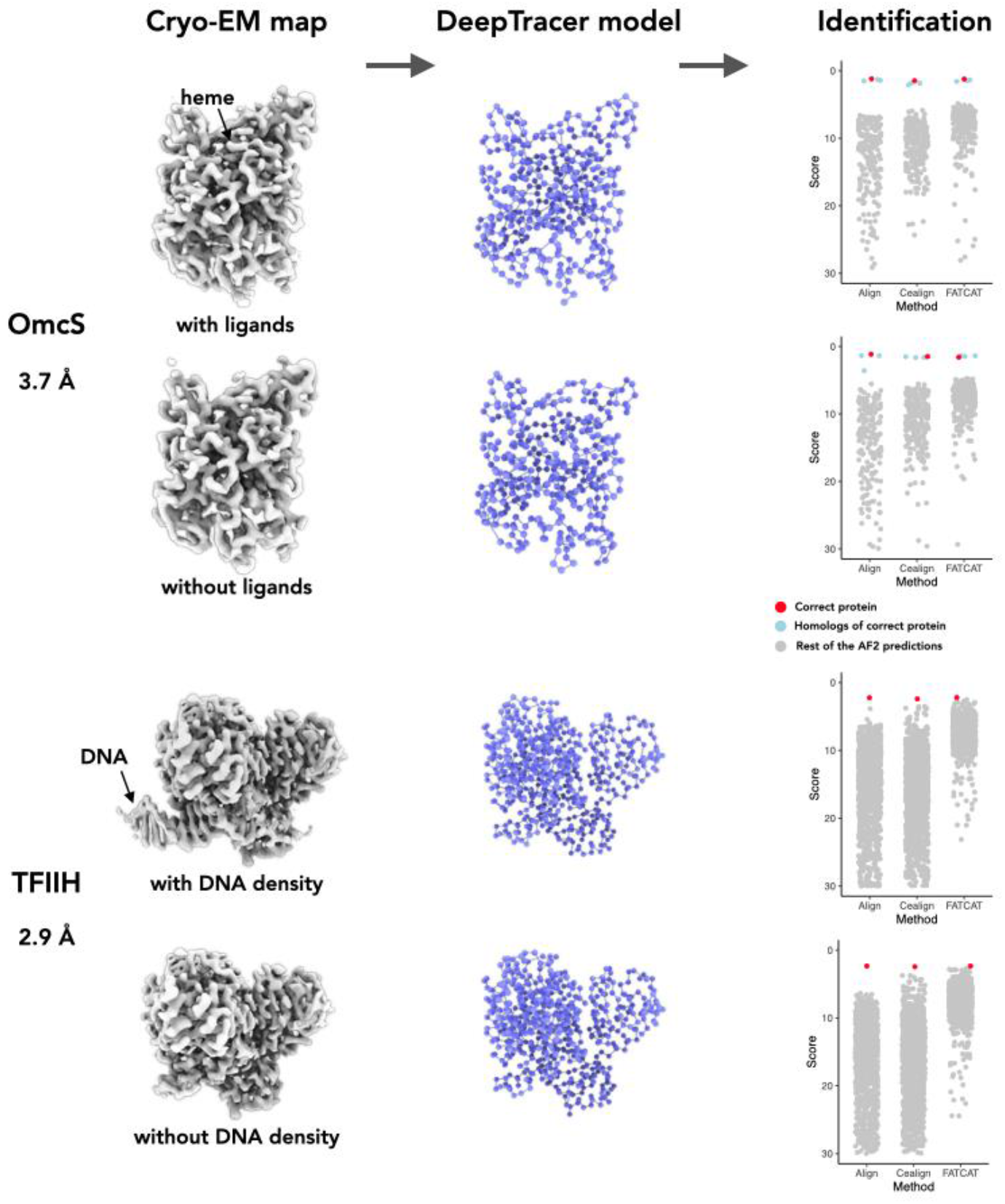
Protein identification is not affected by ligand or nucleic acids densities. Left, cryo-EM maps used for *de novo* protein identification with their reported resolution. OmcS has six hemes per protein subunit. TFIIH protein shown with and without bound dsDNA. Middle, the Cα backbone of the model generated by DeepTracer, from the maps on left. Some extra resides were assigned in the heme area by DeepTracer, while the dsDNA densities are recognized as non-protein area so very few residues were placed there. Right, the DeepTracer-ID scores of AF2 predictions. The correct protein is shown by a red dot, the proteins with significant structural similarity to the correct protein are shown as blue dots, and the remaining AF2 predictions are shown as grey dots. The size of AF2 library and the corresponding organism for the eight benchmark datasets are: OmcS (*G. sulfurreducens PCA*, N=226), and TFIIH (*H. sapiens*, N=1347).

**Fig. S2.**
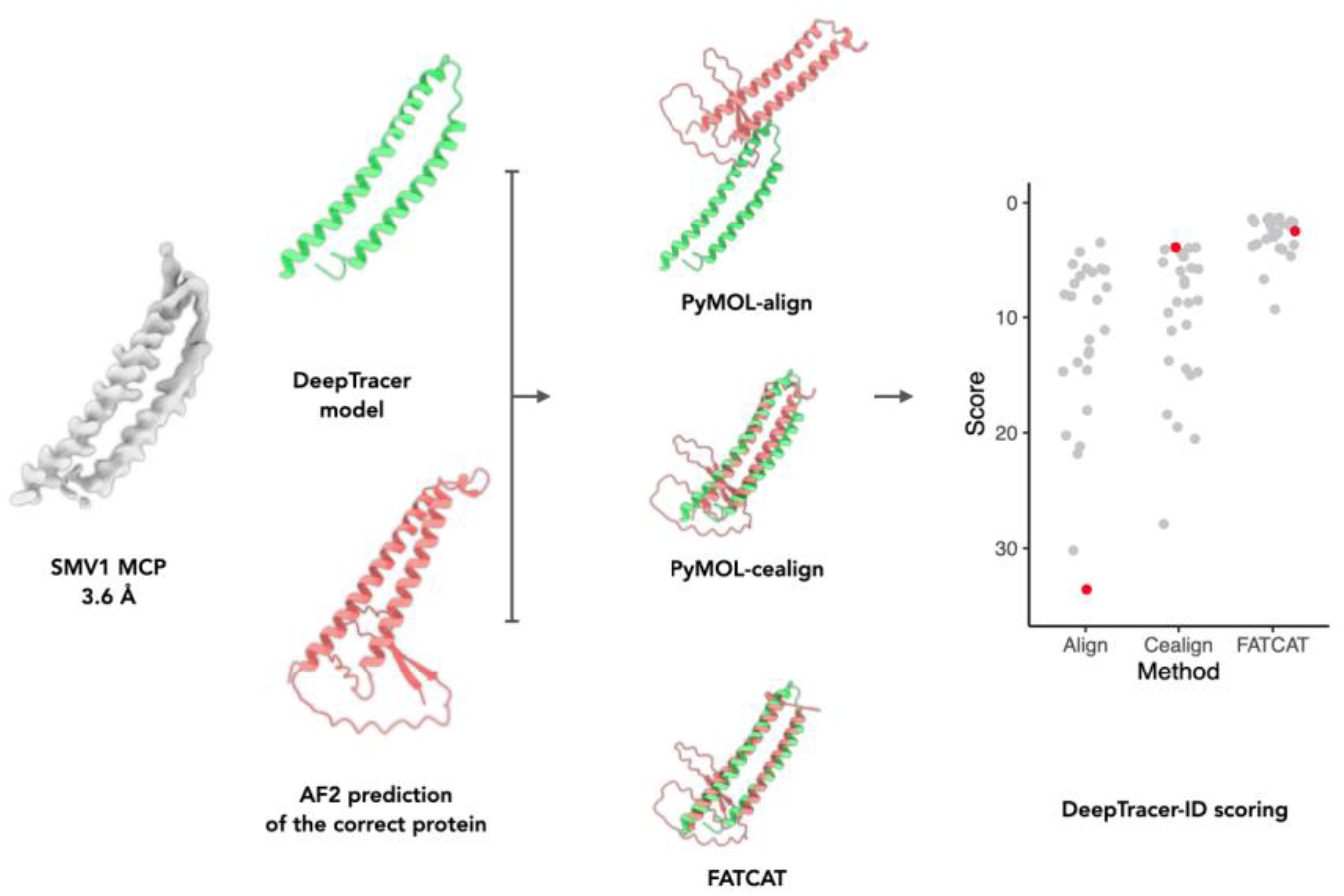
Identifying very small proteins relies on the initial 3D alignments. Far Left, the segmented cryo-EM map of SMV1 major capsid protein with the reported resolution. Left, the DeepTracer model (green) and the AF2 model of the correct protein (red). Right, How the AF2 model is aligned to the DeepTracer model using three different approaches. Far right, the DeepTracer-ID scores of AF2 predictions (SMV1, N=28).

